# Spatially Resolved Determination of Small Molecule–Protein Affinities by Hydrogen–Deuterium Exchange Mass Spectrometry

**DOI:** 10.1101/2025.11.13.688232

**Authors:** De Lin, Luma Godoy Magalhaes, Joel McMillan, Thomas C. Eadsforth, Greg Stewart, Kieran R. Cartmill, Vincent L. G. Postis, Glenn R. Masson

## Abstract

Hydrogen Deuterium Exchange Mass Spectrometry (HDX-MS) is an established tool in drug discovery, used to characterize target engagement and conformational dynamics, frequently used in both biopharmaceutical and small molecule development. Conventional HDX-MS experiments are performed at saturating ligand concentrations to generate a binding “footprint”, where decreased solvent exchange reflects a local structural stabilization and/or reduced solvent accessibility upon binding. Here, we present an extended HDX-MS and HDX-MS/MS titration workflow with electron capture dissociation (ECD) fragmentation capable of estimating apparent dissociation constants (K_D_^app^) at global, peptide and single amino acid resolution by fitting uptake-concentration relationships under EX2 exchange and Langmuir binding assumptions. The ability to determine affinity constants in a spatially resolved manner combined with the automation available in HDX-MS sample handling and data analysis enables quantitative mapping of ligand-protein interactions and provides a scalable approach for structure-activity relationship studies in drug discovery.

## Introduction

Hydrogen Deuterium Exchange Mass Spectrometry (HDX-MS) has emerged as a key technique in Structure Based Drug Discovery, enabling a rapid characterization of protein-ligand interactions and conformational dynamics in both biopharmaceutical and small molecule development (*1–8*). Typically, HDX-MS studies follow a “bottom-up” workflow in which the therapeutical target is incubated with a saturating concentration of ligand in the presence of a deuterated buffer for a series of defined timepoints. Hydrogen-deuterium exchange is then quenched and the protein chemically denatured. The denatured protein is then proteolytically digested, and the resulting peptides are analyzed by LC-MS to determine the extent of deuterium incorporation across their sequences (*9*). The resulting data provide peptide-level special resolution, typically 5-35 amino acids, allowing the identification of regions stabilized or shielded for the solvent upon ligand binding (*10*). Single-residue resolution can be achieved computationally when high peptide redundancy and sequence coverage is obtained (*11, 12*).

To minimize heterogeneous exchange behavior, ligand-binding HDX-MS experiments are commonly conducted at concentrations well above the compound dissociation constant (*K_D_*), producing a fully bound complex in order to avoid heterogenous bi-modal isotopic envelopes (*2*). Recently there has been renewed interest, notably from the Wilson group and co-workers, in HDX-MS/MS (*i.e.* the fragmentation a deuterated precursor peptide to product deuterated product ions) which enables residues-level investigation of deuterated peptides (*13, 14*). These experiments build on the comprehensive and groundbreaking work in the groups of Jørgensen and Rand, who used electron capture dissociation (ECD) or electron transfer dissociation (ETD) to achieve fragmentation with minimal hydrogen/deuterium (H/D) scrambling (*15–18*). Although HDX-MS/MS has been applied to ligand binding (*19, 20*), its use has remained limited, possibly due to the low efficiency of fragmentation and the stringent controls required to detect signal scrambling. Newer technologies, such as Electron-Activated Dissociation (EAD), have shown an improved performance, increasing the accessibility of scramble-free HDX-MS/MS for ligand screening (*13*).

Here, we report a compound titration experiment using both HDX-MS and ECD-based HDX-MS/MS that enables the determination of apparent dissociation constants (K_D_^app^) with spatial resolution. Using the *Plasmodium falciparum* GCN5 bromodomain as a model system, we measured deuterium uptake as a function of compound concentration at the peptide and single-amino-acid level to extract quantitative affinity information. Comparison with orthogonal biophysical methods showed strong agreement with HDX-derived and reference K_D_ values. This approach provides a time efficient, low sample consumption, in-solution method of quantifying ligand binding and mapping local conformational changes, enabling detailed characterization of mode of interaction and binding mechanisms in protein-ligand events.

## Methods

### Materials

Deuterium Oxide (99.9%) and formic acid were purchased from Sigma-Aldrich (St Louis, MO, USA). Ultra-pure HPLC grade water and Acetonitrile were purchased from VWR Chemicals (Radnor, PA, USA). Sodium iodide (2 µg/µL in 50:50 isopropanol:water), 200 pg/nL leucine enkephalin (50:50 acetonitrile:water, 0.1% formic acid), ACQUITY UPLC BEH C18 columns (130Å, 1.7 µm), Enzymate Protein Pepsin Column (300Å, 5 µm, 2.1 mm X 30 mm), glass sample vials (12 x 32 mm screw neck vial, total recovery, 1 mL volume with cap and preslit PTFE/Silicone Septum), as well as all fluidics and trap components were obtained from Waters Corp. (Framingham, MA, USA).

#### Protein expression and purification

The *Plasmodium falciparum* GCN5 bromodomain (residues 1356–1460) was synthesized with an N-terminal TEV-cleavable His₆ tag (Twist Bioscience) and cloned into a pET29a vector using *Nde*I/*Xho*I restriction sites. The plasmid was transformed into *E. coli* BL21(DE3) cells and expressed in Terrific Broth supplemented with 50 µg mL⁻¹ kanamycin. Cultures were grown at 37°C to an OD₆₀₀ of 0.6–0.8, induced with 0.15 mM IPTG, and incubated at 20°C for 18 h. Cells were harvested and lysed in 20 mM HEPES, 500 mM NaCl, 10 mM imidazole, 1 mM TCEP, 5% (w/v) glycerol, pH 7.5, containing DNase I and protease inhibitors cocktail (Merck/Roche, cat. 04693132001), using a continuous-flow cell disruptor (Constant Systems, 30 kpsi).

The His₆-PfGCN5 bromodomain used for Surface Plasmon Resonance experiments was purified by Ni–NTA affinity chromatography (10–500 mM imidazole gradient elution) followed by size-exclusion chromatography (Superdex 75, Cytiva).

For HDX-MS and HDX-MS/MS studies, the His₆ tag was removed by TEV protease digestion after the initial Ni–NTA purification. The cleaved protein was subjected to reverse Ni–NTA purification and further purified by size-exclusion chromatography (Superdex 75, Cytiva). All protein preparations were concentrated to 10 mg mL⁻¹ in 10 mM HEPES, 100 mM NaCl, 0.5 mM TCEP, pH 7.5, flash-frozen, and stored at –80°C until use.

#### Surface Plasmon Resonance (SPR)

Surface plasmon resonance (SPR) experiments were performed on a Biacore™ 8K+ instrument (Cytiva). Flow cell 1 (FC1) served as the reference surface. His_6_PfGCN5 (8 ug/mL) was immobilised on flow cell 2 FC2 of either a NTA derivatized linear polycarboxylate hydrogel (medium charge density, 1500 M chip Xantec, cat n. SCBS NiHC1500M) or a Biacore Sensor Chip NTA (Cytiva, cat. BR100532) using nickel capture followed by amine coupling (amine coupling kit, Cytiva cat. BR100050). Two channels were used in parallel, yielding typical immobilization levels of approximately 2500 RU (channel 1) and 3000 RU (channel 2) was obtained for PfGCN5. The running and dilution buffer contained 25 mM HEPES adjusted to pH 7.5, 150 mM NaCl, 0.5 mM TCEP.

Compounds were prepared as a six-point, three-fold serial dilution (top concentration 30 µM, 0.3% DMSO) using an echo acoustic dispenser (Beckman) to pre stamp a 384-well plate (Greiner, cat. 78128) followed by the addition of 100 µl of dilution buffer to well.

Blank buffer injections (25 mM HEPES pH 7.5, 150 mM NaCl, 0.5 mM TCEP, 0.3% DMSO) were included between compound injection to enable referencing (FC2- FC1 subtraction, followed by compound - blank cycle subtraction). Samples were injected at 30 µL/min over both FC1 and FC2, with a 60 seconds association phase and a 120 seconds dissociation phase. Solvent correct was applied using a five-point DMSO calibration series (0.1-0.5%). The control compound (GSK4027(*23*)) was analysed at the start and end of each run as a six-point, three-fold dilution (top concentration 10 µM) in both channels. Sensorgrams were processed and fitted using Biacore™ Insight Evaluation Software v5.0.18.22102. Equilibrium dissociation constants (K_D_) were determined from steady-state binding responses.

#### Hydrogen Deuterium Exchange-Mass Spectrometry

Prior to deuterium exchange experiments, a protein sequence coverage map was generated by analysing undeuterated PfGCN5 protein under the following conditions: 60 µL of the 1 µM protein in 20 mM HEPES, pH 7.5, 150 mM NaCl, 0.5 mM TCEP and 1% (v:v) DMSO was mixed with to 50 µL ice-cold quenching buffer (4 M Urea, 2% formic acid).

Stocks solution of PfGCN5 20 µM and PfGCN5 20 µM preincubated for 5 minutes at 20°C with 100 µM of compound (DDD0244420) were prepared in 20 mM HEPES pH 7.5, 150 mM NaCl, 0.5 mM TCEP and 1% (v/v) DMSO and maintained at 0.1°C. These mixtures were equilibrated for 5 minutes at 20°C prior to deuterium exchange.

The exchange reaction was performed at 0, 30, 300 and 3000 seconds using the automated liquid-handling platform. For each labelling reaction, 3 µL of sample stock was mixed with 57 µL of D_2_O buffer (20 mM HEPES pH 7.5, 150 mM NaCl, 0.5 mM TCEP and 1% DMSO in 94% D_2_O) at 20°C resulting in a final D_2_O content of 89.4% (v:v).

After the labeling, 50 µL of each sample was mixed with 50 µL of 0.1°C quenching buffer at 0.1°C for 1 minute prior to LCMS data acquisition. The two shortest labelling time points (0.3 second and 3 seconds) were prepared manually, as these durations were below the minimum operating cycle of the liquid-handling system. For the 0.3 seconds exchange time point, the samples were prepared on ice in a 4°C room with pre-chilled D_2_O buffer and pipette tips. For the 3 seconds time point, the samples were prepared and incubated with D_2_O buffer for 3 seconds at room temperature. Samples were frozen immediately into liquid nitrogen and later stored at-80°C until further analysis. All deuterium-labelled samples were repeated three times.

For affinity determination, the experiment was conducted by diluting 3 µL of the ligand-free protein stocks (20 µM) with the 57 µL of D_2_O buffer containing various compound concentrations of 1, 2, 4, 8, 16, 32, 64, and 128 µM while keeping the total protein concentration constant. These mixtures were equilibrated for 5 minutes at 20°C prior to deuterium exchange. The labelled sample (50 µL) was mixed with 50 µL of quenching buffer (4M Urea, 2% formic acid) at 0.1°C for 1 min prior to LCMS data acquisition.

HDX-MS Data were acquired using the HDX manager (Waters Corp.) maintained at 0.1°C and coupled in-line to a SELECT SERIES Cyclic IMS QTOF mass spectrometer (Waters Corps). Automated liquid handling was performed using a PAL3 robotic Tool Change system controlled by Chronos Software (LEAP Technologies). The protein was digested in-line for 2 minutes at 20 °C using an Enzymate BEH Pepsin 2.1 x 30mm Column (Waters Corp.). The generated peptides were desalted in buffer A (0.1% formic acid in water) using an ACQUITY UPLC BEH C18 VanGuard Precolumn (1.7 µm, 5 mm × 2.1 mm; Waters Corp.) prior to reverse-phase separation on an ACQUITY UPLC C18 column (1.7 µm, 100 mm × 1 mm; Waters Corp.). Peptides were eluted using a 40 uL/min flow rate from 5% to 85% buffer B (0.1% formic acid in acetonitrile) in buffer A. Data were acquired in positive mode over a m/z range of 50-1200 with a spray voltage of 2.0 kV. Peptide identification and data processing were performed using ProteinLynx Global Server (PLGS v3.0.3 Waters Corp.). A reference peptide map was generated from three replicates of non-deuterated samples acquired in HDMSe, using identical chromatographic and instrumental parameters to those applied for the deuterated samples.

Raw data were processed using HDExaminer v3.5.0 (Sierra Analytics) or Newmarket (unreleased software, Waters Corp.) for deuterium uptake analysis. Reported uptake values were not corrected for back-exchange and therefore represent relative deuterium incorporation. All mass spectrometry data have been deposited to the ProteomeXchange Consortium via the PRIDE partner repository with the dataset identifier PXD070051.

Equilibrium dissociation constants (K_D_^app^) were obtained from steady-state binding response by fitting ΔD versus ligand concentration to a Langmuir isotherm (1:1) under EX2 exchange, assuming rapid conformational pre-equilibrium and negligible ligand depletion.

#### Derivation of local protein ligand dissociation constant from HDX-MS data

Differences in deuterium uptake as a function of ligand concentration were used to monitor local protein–ligand interactions at peptide-or single-amino-acid-level spatial resolution. The percentage of deuterium uptake protection (ΔD) was calculated by comparing uptake in the presence and absence of ligand at 300 seconds of exchange reaction at 20°C. Data exported from HDExaminer v3.5.0 (Trajan Scientific and Medical) or Newmarket (beta, unreleased software; Waters Corp.) were analyzed using a custom automated Python script (HDX KD data analysis HDXaminer.py or ECD_KD_all_peptide_final.py respectively). The script parses peptide-level uptake values, aligns data across conditions, calculates differential deuterium uptake (ΔD), and generates comparative plots of HDX kinetics. The Python analysis scripts are available at Zenodo (DOI: 10.5281/zenodo.17492311) and maintained on GitHub (https://github.com/Glenn-Masson/De_HDX_KD).

### Targeted ECD

Peptides exhibiting protection upon compound binding in HDX-MS experiments were selected for targeted electron capture dissociation (ECD) fragmentation based on exact mass and retention time and window. ECD cell parameters were optimized to minimize hydrogen/deuterium scrambling using the “P1 peptide” standard (sequence HHHHHHIIKIIK, synthesised by Cambridge Bioscience) (*24*) and by monitoring deuterium lass coincident with ammonia ion fragment formation (*17*). ECD fragment spectra were processed using Newmarket software (Beta, unreleased software, Waters Corp.), visualized with SigmaPlot v14.5 (Systat Software) and PyMOL v3.0.0 (Schrödinger LLC).

## Results and Discussion

### Dissociation Constants Determined via Peptide Level HDX-MS Analysis

To demonstrate the capacity of HDX-MS analysis to determine dissociation constants for ligand binding, we used the bromodomain (BRD) of *Plasmodium falciparum* histone acetyltransferase GCN5 (General Control Nonderepressible 5) (PfGCN5) (residues 1356-1460). PfGCN5-BRD is a potential target for the treatment of malaria parasite *Plasmodium falciparum* (*25*), which has a classical bromodomain structure of four alpha helices with two interhelical loops (*21, 22, 26*). An in-house drug discovery screen developed the compound DDD02444250 (‘4250), which, using surface plasmon resonance (SPR) had a reported K_D_ of 4.8 μM (supplementary figure 1).

To determine the site of interaction of ‘4250 with PfGCN5-BRD, we conducted HDX-MS using a Trajan LEAP Pal autosampler, with a Waters HDX-Manager upstream of a Waters cIMS instrument, fitted with an eMision Electron Capture Dissociation (ECD) cell. We were able to create a peptide map of PfGCN5-BRD consisting of 84 peptides, representing 100% coverage with a 10.17 redundancy score (Figure 1A). Differential HDX-MS was performed by incubating 20 μM of PfGCN5-BRD with 20 μM ‘4250 prior to deuteration at five different timepoints (0.3s, 3s, 30s, 300s, and 3000s – with the 0.3s timepoint being a 3 second timepoint conducted on ice) to produce a deuterium uptake difference map between the apo and ligand bound states (Figure 1B,C,D and Table 1.0).

**Figure 1:**
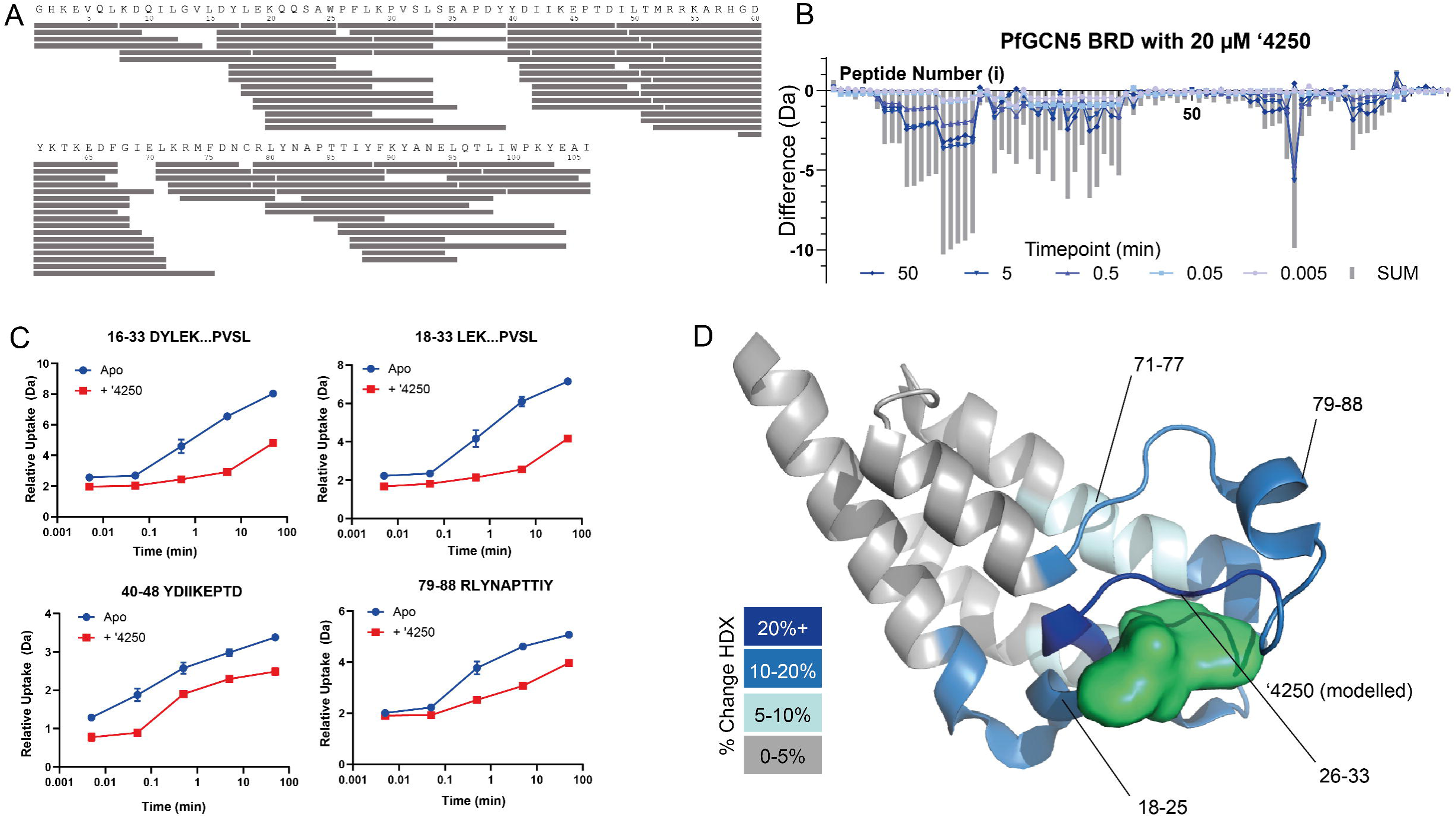
Global shifts in solvent exchange in PfGCN5 on compound binding. (A) Peptide map of PfGCN5BRD showing coverage of pepsin-produced peptide map. (B) Global HDX profile of PfGCN5-BRD on incubation with 20 µM ‘4250. Each timepoint is shown alongside the sum of the shifts (SUM). (C) Representative peptide uptake plots of four peptides showing changes on’4250 binding. Each data point is the average with standard deviation shown as error bars (n=3) (D) Shifts in HDX mapped onto PfGCN5BRD crystal structure PDB: 5TPX. The compound ‘4250 is modelled into the existing compound binding site.

**Table 1.0:** HDX-Statistics Table for ‘4250 binding to PfGCN5-BRD. * = 0.3s second timepoint conducted as a 3 second timepoint on ice. ** = conducted using the LEAP liquid handling robot.

|  |  |  |
| --- | --- | --- |
|  | Apo | 4250 |
| # of peptides | 82 | 82 |
| Coverage | 100% | 100% |
| Average Peptide Length | 12.59 aa |  |
| Redundancy | 9.75 |  |
| Timepoints | 0.3s*, 3s, 30s**, 300s**, 3000s ** |  |
| Replicates | ND 2, Deuterated 3 |  |
| Replicability | 0.89% |  |
| Significance Threshold | >5%/0.5 Da/ Passes t-test |  |

Next, we sought to determine whether the apparent dissociation constants (K_D_^app^) of the compound ‘4250 for PfGCN5-BRD could be quantified by HDX-MS titration. An 8-point 1:1 dilution series of ‘4250 was prepared from a starting concentration of 120 μM in the presence of 20 µM PfGCN5-BRD and deuterium exchange was performed at a single 5 minutes timepoint (Figure 2). This labeling period was chosen to maximize the dynamic range of measurable protection, with all peptides exhibiting the largest differences in deuterium uptake upon compound binding, whereas differences in H/D exchange kinetics converged at shorter and longer labeling intervals.

**Figure 2:**
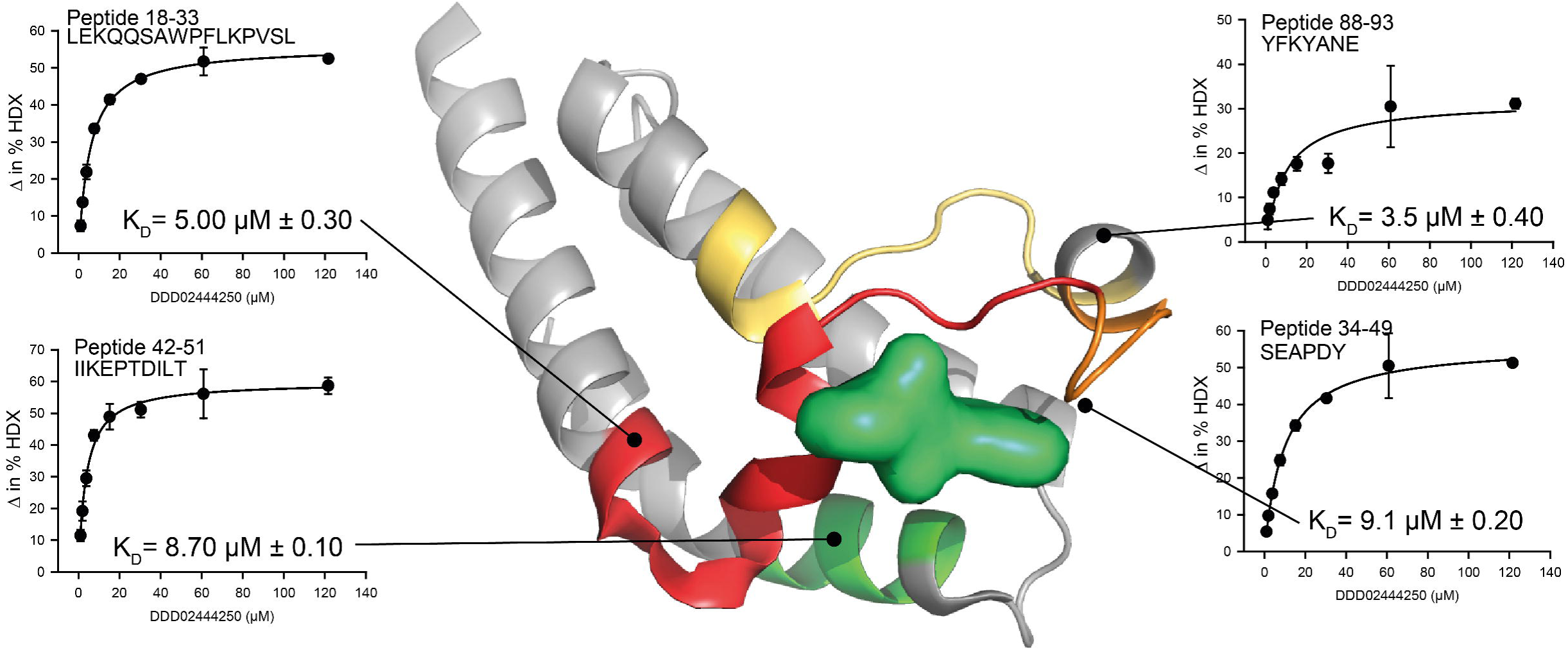
Peptide Level HDX-MS to derive spatial K_D_^app^. In total, the K_D_^app^ values for over 50 peptides could be determined (see Table 2.0). Each data point is the average of three independent exchange reactions with standard deviation shown (n=3).

**Table 2.0:**
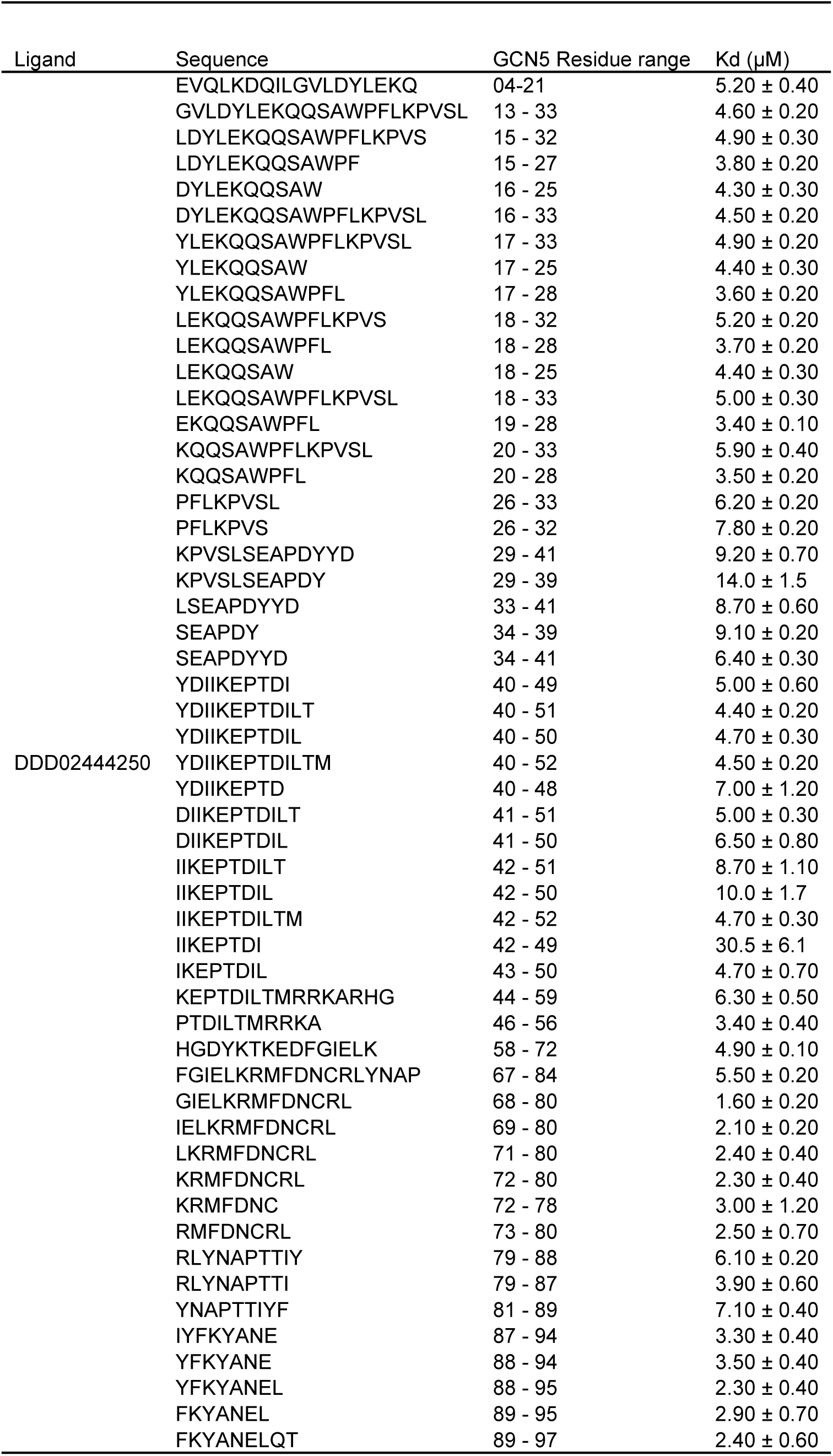
Dissociation Constants for peptides exhibiting a shift on ‘4250 incubation. Each Kd value is derived from three independent exchange reactions (n=3).

ΔD values representing the change in deuterium uptake between apo and ligand-bound PfGCN5-BRD were obtained at the 5 minutes labeling time point each compound concentration in the titration series. Apparent dissociation constants (K_D_^app^) values were determined by fitting plots of ΔD versus ligand concentration to a single-site Langmuir equation:

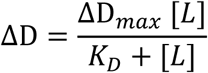

where ΔD_max_ is the fitted maximal deuterium uptake protection, [L] is the ligand concentration, and Kd is the fitted local equilibrium dissociation constant. Non-linear regression was performed using a custom python script (HDX KD data analysis HDXaminer.py), and the coefficient of variation (CV) of ΔDₘₐₓ was calculated using the standard deviation from curve fitting to assess fitting precision. A selection criterion was applied such that a Kd value was accepted only if ΔDₘₐₓ was at least 20% and its CV was below 20%.

The resulting titration data yielded characteristic binding curves showing a plateau in deuterium exchange protection as a function of compound concentration. From these fits, the midpoint of each curve corresponded to the apparent dissociation constant (K_D_^app^). A total of 52 peptides produced measurable K_D_^app^ values for compound ‘4250, many representing overlapping sequence regions of PfGCN5-BRD, providing high internal consistency (Table 2). The mean (K_D_^app^) was 5.5 ±4.1 μM, in close agreement with the reference value obtained by SPR (4.8 μM Cl=4.0-5.8 95%). Individual K_D_^app^ peptide-derived values ranged from from 1.6 ±0.2 μM (residues 68-80) to 30.5 ± 6.1 μM (residues 42-49). These results demonstrate strong concordance between peptide-level HDX-MS-derived affinities and orthogonal biophysical measurements, supporting the reliability of this approach for quantifying ligand binding in solution.

For peptides showing no measurable decrease in deuterium uptake upon compound binding, no K_D_^app^ could be determined. The titration experiment was repeated at the 30s and 3000s labeling times to evaluate the effect of exchange duration on the calculated K_D_^app^. At 3000 s, most deuteration curves converged, yielding no observable ΔD_max_ and therefore preventing K_D_^app^ estimation. Similarly, at 30 s, deuteration rates had not diverged sufficiently between bound and unbound states to provide reliable fitting of the binding curve. These observations highlight the importance of selecting an optimal labeling time that maximizes the dynamic range of measurable protection; future implementation of an automated time-point optimization tool would facilitate rapid identification of conditions most suitable for K_D_^app^ determination across all peptides.

### Dissociation Constants Determined via Peptide Level HDX-MS/MS Analysis

We next sought to determine whether ECD could be used under minimized H/D-scrambling conditions to determine apparent dissociation constants (K_D_^app^) at single-amino-acid resolution. Source and ECD parameters were optimized using the P1 peptide HDX scrambling monitor to tune as previously described (*16, 19, 24, 27*) to ensure minimal deuterium redistribution. Targeted ECD fragmentation was then applied to deuterated precursor ions of the PfGCN5-BRD peptide containing residues 18 to 33 which covers the tightest region of binding. This peptide, detected predominantly in +4 charge state and with a relative abundance exciding 10^5^ counts, produced an almost complete *c* ion series including *c*3-*c*7, *c*9-*c*11 and *z* ion fragments *z*9-*z*13, providing a near-complete single-residue coverage (Figure 3.0).

**Figure 3:**
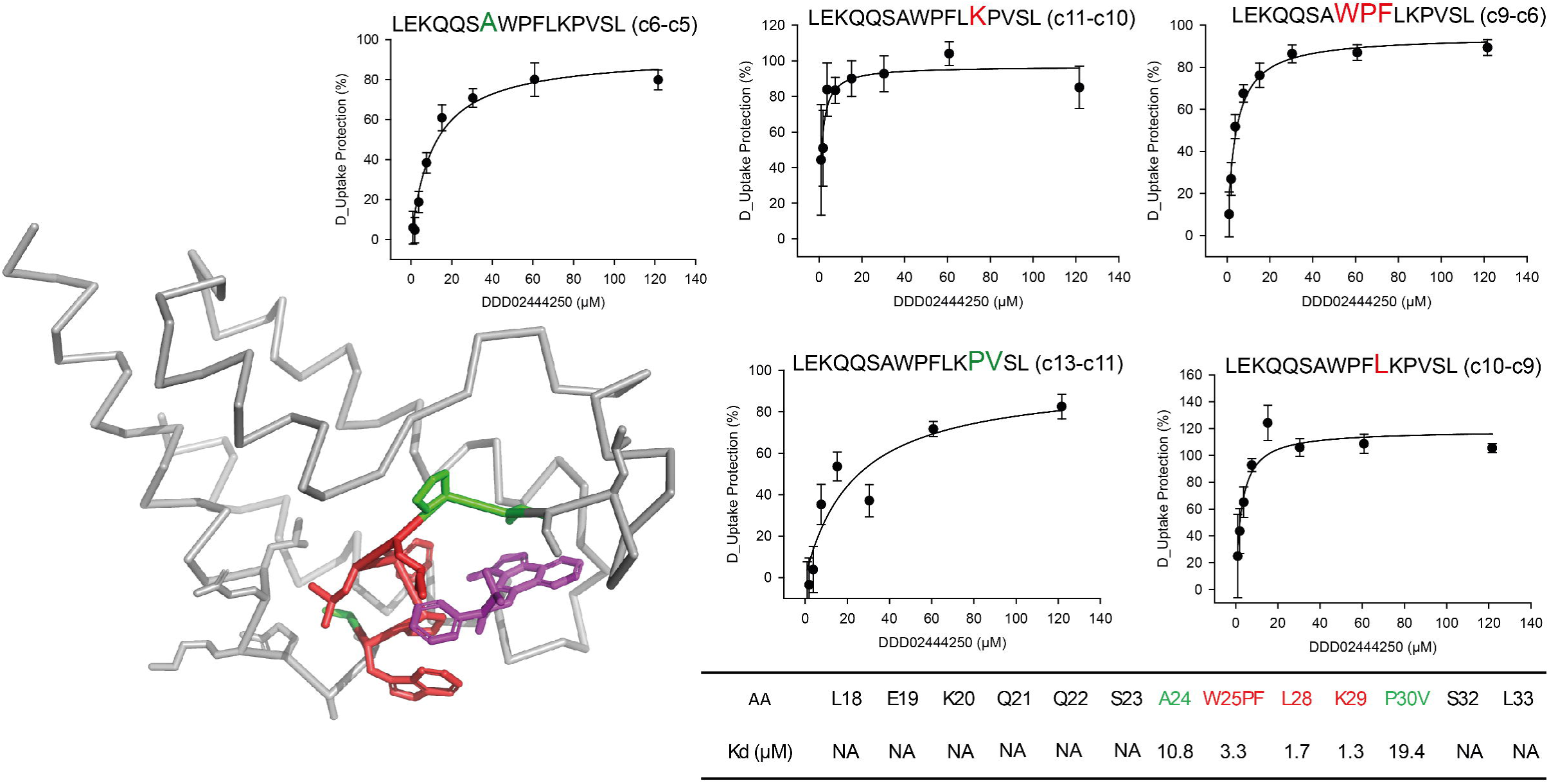
Single Amino-Acid HDX-MS to derive spatial K_Ds_ at a single residue resolution. HDX-MS/MS was conducted using targeted ECD on a single peptide while titrating the ‘4250 compound. Consecutive fragments are used to determine individual exchange rates, from which % deuteration can be derived. Each data point is average with standard deviation, from n=3, with each exchange reaction being conducted independently.

Compound titration of ‘4250 across eight concentration curves enabled the determination of residue-level K_D_^app^ values for the interacting region A24-V31 (Figure 3). The apparent affinities ranged from 19.4 µM for P30V to 1.3 µM for K29. Notably, this stretch of amino acids contains two proline residues, P30 and P25, which are invisible to HDX-MS/MS analysis due to the absence of backbone amide hydrogens. The HDX-MS/MS analysis therefore not only localized the ligand binding interface between PfGCN5-BRD and ‘4250, but also provided a quantitative, residue specific affinity information.

## Conclusion

Recent advances in peptide fragmentation technologies for HDX-MS/MS (*2, 13, 14, 28*) have renewed interest in achieving residue-level localization of deuterium uptake. Electron-activated dissociation (EAD) allows an efficient fragmentation of deuterated peptides while minimizing hydrogen/deuterium (H/D) scrambling. Here, we demonstrate that HDX-MS/MS using ECD can not only localize ligand binding sites but, through compound titration at selected exchange time points, can also be used to derive apparent dissociation constants (K_D_^app^). This represents a novel and generally applicable in-solution approach to quantifying ligand affinity without the need for protein tagging or immobilization (*2, 13, 14, 28*).

A key advantage of HDX-MS/MS is its ability to resolve affinity information at single-amino-acid spatial resolution, providing insight into local modes of interaction that complement traditional Structure–Activity Relationship (SAR) studies. The combination of HDX-MS-derived spatial interaction mapping with conventional SAR analysis can accelerate hit validation and early lead optimization by linking compound structure to localized protein response. The approach is compatible with standard HDX-MS instrumentation and can be readily adapted to other protein–ligand systems, particularly where in-solution binding measurements are advantageous. The affinity measurement can also be conducted without the use of protein affinity tags (such as His/Strep-tags), or the need for fluorescent moieties which may introduce artefacts into the experiment. Future developments in automated ECD data acquisition and analysis are expected to further improve throughput and robustness, expanding the applicability of this approach in drug discovery.

## Acknowledgements

We acknowledge funding from BBRSC (BBSRC Capital Equipment Fund BB/V019635/1) for support for HDX-MS. We gratefully acknowledge financial support from Wellcome Trust Centre Award [203134/Z/16/Z]. GRM is supported by a Baxter Fellowship, University of Dundee. The authors thank Waters for providing early access to prototype data-analysis software that facilitated the ECD data processing, and in particular Dr. Owen Cornwell and Dr. Dale Cooper-Shepherd for their technical support and collaboration.

## Author Contributions

DL: Investigation, Methodology, Conceptualization Resources, Writing—review and editing. LGM, TCE, JMM, GS, KC: Investigation. V.P.: Supervision, Conceptualization, Resources, Writing—review and editing, Project Administration. G.R.M.: Conceptualization, Resources, Supervision, Investigation, Funding acquisition, Methodology, Writing—original draft, Project administration, Writing—review and editing.

## Data Availability

All the mass spectrometry data and HDExaminer files have been deposited to the ProteomeXchange Consortium via the PRIDE partner repository with the dataset identifier PXD070051.

---

**Supplementary Figure 1:**
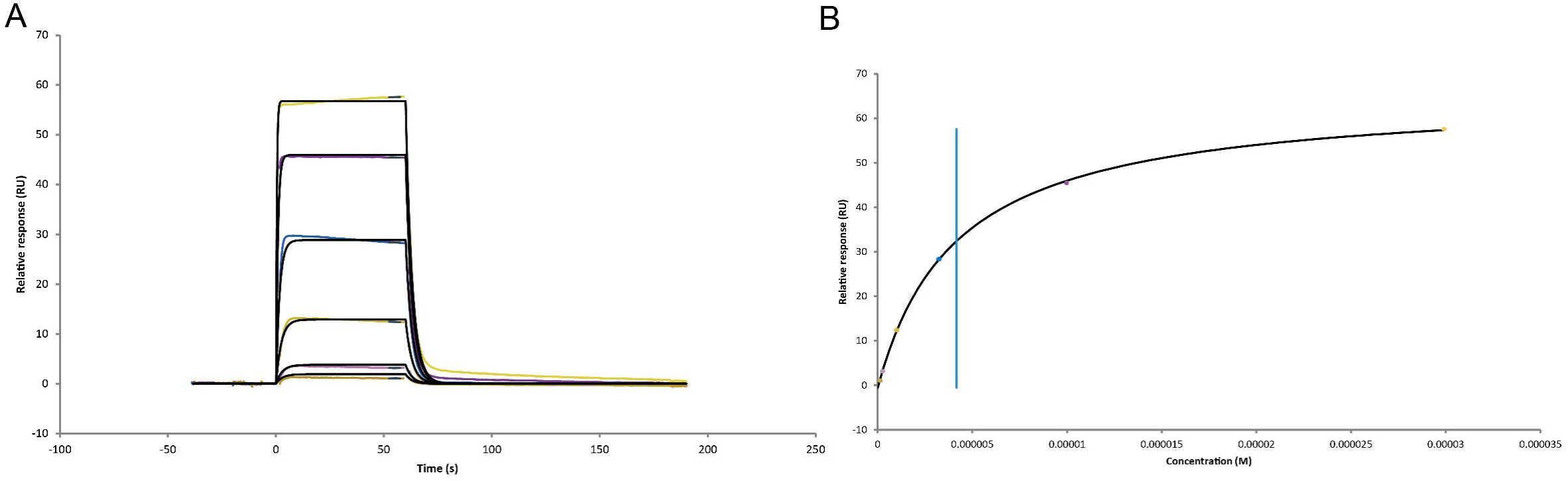
Surface Plasmon Resonance (SPR) determination of dissociation constant of ‘4250 for PfGCN5 BRD. (**A**) A 6-point concentration curve showing 1:1 binding of’4250 to immobilized PfGCN5 BRD. (**B**) Dissociation Constant determination of ‘4250 based on maximal steady state response obtained in (A). Blue line represents mid-point of the curve and associated dissociation constant. Data representative of three independent repeats.

